# Effect of Antimicrobial Peptides HNP-1 and hBD-1 on *Staphylococcus aureus* Clinical Strains *in vitro* and *in vivo*

**DOI:** 10.1101/578278

**Authors:** Albert Bolatchiev, Vladimir Baturin, Igor Bazikov, Alexander Maltsev, Elena Kunitsina

## Abstract

The aims of this study were: (1) To investigate the activity of recombinant AMPs HNP-1 and hBD-1 in combination with cefotaxime against *Staphylococcus aureus* clinical strains (MSSA and MRSA) *in vitro* using checkerboard method; (2) To investigate the activity of HNP-1 and hBD-1 encapsulated in silicon nanoparticles (niosomes) in the treatment of MRSA-infected wound in rats. For this *S. aureus* clinical strains (MSSA and MRSA) were isolated from patients with diabetic foot infection. Cefotaxime, recombinant HNP-1 and hBD-1 (in all possible combinations with each other) were used for testing by the checkerboard method. Two niosomal topical gels with HNP-1/hBD-1 were prepared to treat MRSA-infected wounds in rats. Gels were administered once a day, the control group – without treatment. Wound healing rate was calculated on the 4th, 9th and 16th days of the experiment and compared using one-way ANOVA with Bonferroni correction. MIC of HNP-1 for MSSA and MRSA was the same – 1 mg/L. MIC of hBD-1 for MSSA and MRSA was also the same – 0.5 mg/L. Topical gels with niosomal HNP-1 (or hBD-1) showed a significantly faster wound healing in comparison with the control. The data obtained open up prospects for use of AMPs encapsulated in silica nanoparticles for the development of new antibiotics.

## INTRODUCTION

The human body is in a constant contact with various pathogens and the evolution has developed protective mechanisms that can prevent the initiation of infection process at the first contact of an infectious agent with the macroorganism. The adaptive immune response is not able to promptly carry out adequate elimination of pathogens, so the evolutionarily ancient innate immune response comes to its aid. The innate immune system recognizes a wide range of molecular patterns inherent in a huge number of pathogens (PAMP – pathogen-associated molecular pattern) and starts the subsequent stages of the immune response [1].

Currently, clinical medicine faces a huge problem – there is a widespread rapid increase in the resistance of microorganisms to antibacterial drugs [2]. Declining susceptibility of microbes to antibiotics is a natural evolutionary process of pathogen’s adaptation to selective pressure of antibacterials [3].

Thus, it is obvious that medicine in the coming decades will face difficulties in treating infections caused by resistant pathogens, which in turn will lead to an increase in mortality, treatment duration and cost. To combat the growing antibiotic resistance, development of antibacterial drugs is required, the resistance to which does not form over time. These drugs should also have a high efficacy and safety. From this point of view, the development of new antibiotics based on antimicrobial peptides (AMPs which are an important module of innate immunity) is the most promising [4,5]. Among AMPs, the most interesting are human neutrophil peptide-1 (HNP-1, or α-defensin-1) and human β-defensin-1 (hBD-1). These defensins have a wide spectrum of antimicrobial activity against gram-positive and gram-negative bacteria, viruses and fungi. Antimicrobial mechanism of action of positively charged AMPs is based on membrane (negatively charged) permeabilization followed by microbial lysis [4]. One of the problems with AMP’s clinical application is their rapid degradation that leads to a short period of action [6]. To solve this problem, we suggest using the method of encapsulation of HNP-1/hBD-1 in silica nanoparticles (niosomes) [7].

To explore the prospects for a possible clinical use of HNP-1 and hBD-1, we aimed: (1) to investigate the antistaphylococcal activity of HNP-1 and hBD-1 and the synergistic effect with cefotaxime *in vitro*; (2) to study the regenerative activity of topical niosomal gels with recombinant HNP-1/hBD-1 on *S. aureus*-infected wound in rats. *S. aureus* has been chosen as one of the most clinically relevant microorganisms [8].

## MATERIALS AND METHODS

### Bacteria and antimicrobials

*S. aureus* clinical strains (methicillin-sensitive – MSSA and methicillin-resistant – MRSA) were isolated from hospitalized patients with diabetic foot infection. Microflora isolation from the wound discharge was conducted before the antibiotic therapy was initiated. Identification of microorganisms and their antibiotic resistance were carried out in accordance with EUCAST protocols [9] in The Department of Clinical Microbiology of The Center of Clinical Pharmacology and Pharmacotherapy, LLC (Stavropol, Russia).

Recombinant HNP-1 (purity ≥ 92%) and hBD-1 (purity ≥ 95%) (Cloud-Clone Corp., USA; both peptides were expressed in *E. coli*) were used for checkerboard assays and preparations of niosomal topical gels for *in vivo* experiments. Amino acid sequences for HNP-1: EPLQARADEVAAAPEQIAADIPEVVVSLAWDESLAPKHPGSRKNMACYCRIPACIAGERRYGTCIYQGRLWAFCC and for hBD-1: GNFLTGLGHRSDHYNCVSSGGQCLYSACPIFTKIQGTCYRGKAKCCK. Cefotaxime, powder for solution for injection (Claforan, Aventis Pharma Limited, United Kingdom), was used for testing by the checkerboard method – this antibiotic (3rd generation cephalosporin) is often used in hospitalized patients, acts on MSSA and does not affect MRSA.

### Synergy tests by checkerboard method

Standard checkerboard assays were used for synergy tests [10,11]. Pure cultures of *S. aureus*, MSSA and MRSA (clinical strains isolated from patients with diabetic foot infection), were cultivated on solid nutrient medium (Mannitol Salt Agar, BioMedia LLC, Russia) for 18-24 hours, 37ºC. From a fresh morning culture, a suspension was prepared in a sterile saline solution that corresponded to a turbidity standard of 0.5 according to McFarland – that is, the resulting suspension had an approximate *S. aureus* concentration of 1.5 × 10^8^ CFU/mL. Then, 0.1 mL of this suspension was dissolved in 9.9 mL of 2.1% Mueller-Hinton broth (SIFIN Institut für Immunpräparate und Nährmedien GmbH, Germany) and eventually a solution was obtained – inoculum – containing approximately 5 × 10^5^ CFU/mL, which corresponds to the standard EUCAST protocol [12–13].

Then, 100 µl of inoculum was administered into each well of 96-well microdilution plates (U-bottom shape, single-use polystyrene microdilution plates with a lid for immunological reactions, Medpolimer OJSC, Russia). After that, serial twofold dilutions of each antimicrobial agent (HNP-1 or hBD-1 or cefotaxime, 50 µl per well) were administered into the wells. For greater accuracy of the experiment, a triple control was performed – three wells in each plate contained: 1) control-1 – only 2.1% Mueller-Hinton broth (200 µl per well, without bacteria and without antimicrobial agents); 2) control-2 – only bacterial inoculum (200 µl per well, without antimicrobial agents); 3) control-3 – only antimicrobial agents without inoculum (100 µl of substance A + 100 µl of substance B) at maximal concentrations (see Supplementary data).

For checkerboard assays, the following combinations of antimicrobials were selected: 1) cefotaxime + HNP-1; 2) cefotaxime + hBD-1; 3) HNP-1 + hBD-1. Antimicrobial agents were dissolved in 2.1% Mueller-Hinton broth. In all experiments, cefotaxime had a dilution range from 0 mg/L to 32 mg/L; HNP-1 and hBD-1 from 0 mg/L to 5 mg/L. Experiments with each pair of antimicrobial agents were repeated at least three times. After adding inoculum and antimicrobial agents, the plates with the lids closed (to prevent drying) were incubated in a thermostat overnight at 37ºC. In 18-20 hours, the presence or absence of growth was visually evaluated. MIC is the lowest concentration of an anti-infective agent, at which there was no visible growth of microorganisms [14]. The combined microbicidal effect of two substances (A and B) was assessed by the value of the fractional inhibitory concentration index (FICI) [15,16]: *FICI = (A/MIC A) + (B/MIC B)*, where A and B are such concentrations of antimicrobial agents in their mixture that inhibit the growth of bacteria; MIC A and MIC B, respectively, the minimum inhibitory concentrations of substances A and B (not in combination with each other). Depending on the FICI value obtained, there are three types of mutual influence of the two investigated antimicrobial substances on microorganisms: 1) FICI ≤ 0,5 – synergism of action – substances A and B mutually enhance the microbicidal effect of each other; 2) 0.5 < FICI < 4 – no interaction – the effect of two antimicrobial substances does not depend on the presence of each other; 3) FICI > 4 – antagonism – substances A and B reduce the antimicrobial effect of each other [16].

### Niosomal HNP-1/hBD-1 preparation

For the experiments on animals, two topical gels were obtained: a niosomal gel containing HNP-1 (2 mg/L) and niosomal gel containing hBD-1 (1 mg/L). The concentrations of antimicrobial peptides (two times greater than the MIC) were chosen in accordance with the results of *in vitro* experiments. To form silica nanoparticles (niosomes), we used PEG-12 dimethicone which has amphiphilic properties that allow partitioning of the water-soluble part (polyethylene glycol) to water and the fat-soluble part (dimethicone) to lipids. This structure makes it easy to obtain microcapsules by shaking. For the preparation of topical niosomal gels (with 2 mg/L of HNP-1 or 1 mg/L of hBD-1), we used the standard method previously described by us [17]. It should be noted that to form nanoparticles with sizes of 100-140 nm, the obtained preparation of niosomes with encapsulated HNP-1/hBD-1 was placed in an ultrasonic treatment vessel. Sound mode: frequency — 20 kHz, power — 200 W; ultrasonic processing time intervals — 15, 30 and 45 min [17].

### Ethics

Animal experimental protocols were approved by the Ethics Committee of Stavropol State Medical University (Minutes of the meeting of the Ethics Committee No. 52). Wistar rats (males, weighing 180-200 g, 6 weeks of age) were used. Before the start of the study, the animals were kept for a month at identical conditions (in the vivarium of Stavropol State Medical University, two rats in each cage), with enough food and water (*ad libitum*).

### Animal studies protocol

An experimental model of an infected wound was used to study wound healing activity of topical niosomal gels with HNP-1 and hBD-1. Rats’ hair was removed on the site of the proposed application of the wound defect (day 1 of the experiment); cosmetic cream was used for hair removal (Veet, France). This method of hair removing was chosen due to its atraumatic nature, the lack of the need to use a razor and ease of use. The next day (day 2 of the experiment), the rats were wounded with a punch biopsy tool (Medax EPT8000-00, Italy; diameter – 8 mm). Before wounding, the rats were anaesthetized with ketamine (80 mg/kg) and xylazine (10 mg/kg). Wounds were inoculated with 1 mL of *S. aureus* (MRSA isolated from a patient with diabetic foot infection) with a turbidity of 0.5 McFarland (1.5 × 10^8^ CFU/mL).

To control the rate of wound healing, the Wound Pro software for iOS was used to determine the area (mm^2^) of surface defects. A day after inoculation (day 3 of the experiment), the rats were randomly divided into 3 groups: group 1 – control, without treatment; group 2 – niosomal gel with 2 mg/L of HNP-1; group 3 – niosomal gel with 1 mg/L of hBD-1. In each group there were 10 rats (n=10); two rats in each cage. Treatment was initiated two days after injury and infection (day 4 of the experiment). Antimicrobial substances were externally applied once a day to the affected area. The total duration of the treatment was 12 days. The wound area was measured on the 4^th^, 9^th^ and 16^th^ days of the experiment. To assess wound healing, the following formula was used: *w* = (*S*_1_ - *S*_*t*_)/*S*_1_ × *100*) [18], where *w* is the wound healing rate; *S*_1_ is the initial area of the wound (in the beginning of treatment – on the day 4 of the experiment); *S*_*t*_ – the area of the wound after a period *t –* on the 9^th^/16^th^ day of the experiment. The *w* value (wound healing rate, %) was calculated in each group from 4^th^ to 9^th^ day of the experiment (*w*_*4-9*_) and from 9^th^ to 16^th^ day (*w*_*9−16*_).

### Statistical analysis

To compare the wound healing rate differences between three groups, one-way ANOVA test with the Bonferroni correction was used (MaxStat Software, Germany). p<0.05 was considered statistically significant.

## Results

### Antistaphylococcal activity of recombinant HNP-1 and hBD-1 in vitro

The studied antimicrobial peptides were shown to have pronounced antimicrobial activity against both MSSA and MRSA clinical strains isolated from wound discharge of patients with diabetic foot syndrome.

The HNP-1 MIC for MSSA and MRSA was the same – 1 mg/L. The MIC of β-defensin-1 for MSSA and MRSA was also identical – 0.5 mg/L. Furthermore, the cefotaxime MIC for MSSA was determined to be 2 mg/L. The MIC of this antibiotic for MRSA was not determined, since methicillin-resistant strains are not sensitive to β-lactam antibiotics (except fifth generation cephalosporins).

The following combinations of antimicrobial agents against MSSA were investigated: 1) HNP-1 + cefotaxime; 2) hBD-1 + cefotaxime and 3) HNP-1 + hBD-1. In all cases (with experiments done in triplicate), FICI was 1, which indicated no interaction between these defensins and β-lactam antibiotic and no interaction between HNP-1 and hBD-1.

As for MRSA, it is impossible to calculate FICI for the combinations of HNP-1 + cefotaxime and hBD-1 + cefotaxime, since the MIC of cefotaxime for this strain of staphylococci cannot be determined due to its natural resistance. FICI for MRSA with the combined use of HNP-1 and hBD-1 was 1.5, which indicates that these AMPs do not affect each other’s antimicrobial activity. Results of synergy tests by checkerboard method are available as Supplementary data.

### Effect of HNP-1/hBD-1 encapsulated in silica nanoparticles on MRSA-infected wound healing in rats

In group 1 (control, without treatment), the average area of wounds on the 4^th^ day of the experiment was 5.7 ± 0.7 mm^2^, on the 9^th^ day – 5.1 ± 1.4 mm^2^, on the 16^th^ day – 4.2 ± 1.8 mm^2^. In group 2 (niosomal HNP-1, 2 mg/L), the average area of wounds on the 4^th^ day of the experiment was 5.8 ± 1.7 mm^2^, on the 9^th^ day – 2.5 ± 1.3 mm^2^, on the 16^th^ day – 0,6 ± 1.4 mm^2^. In group 3 (niosomal hBD-1, 1 mg/L), the average area of wounds on the 4^th^ day of the experiment was 6.4 ± 1.2 mm^2^, on the 9^th^ day — 3.9 ± 1.5 mm^2^, on the 16^th^ day — 1,2 ± 1.0 mm^2^. Figure 1 presents examples of wound healing in different groups.

**Fig. 1.**
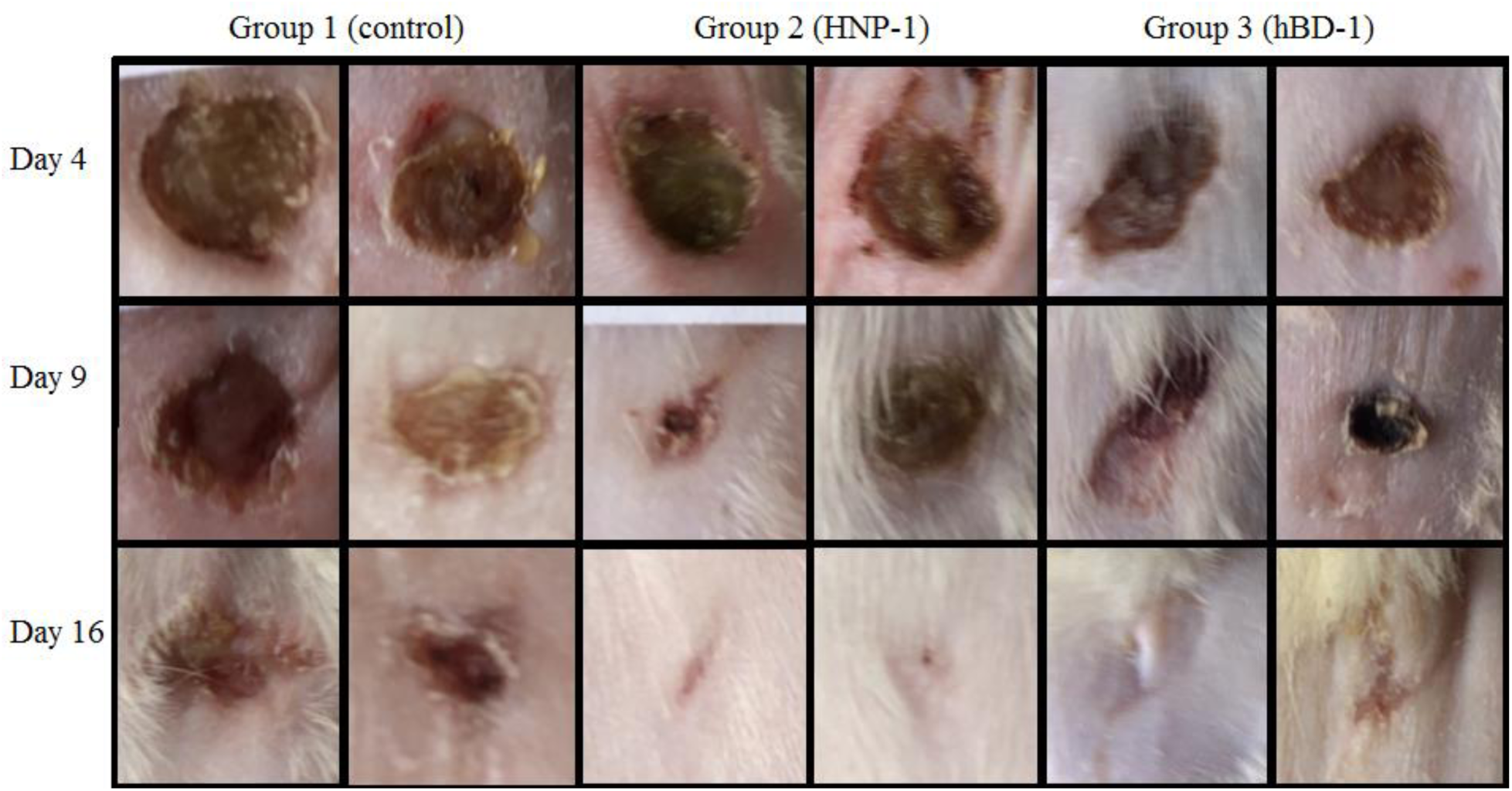
Examples of wound healing in different groups on the 4^th^, 9^th^ and 16^th^ day of the experiment (2 examples from each group are presented).

In group 1, the wound healing rate from the 4^th^ to 9^th^ day of the experiment (*w*_*4-9*_) was 11.0 ± 19.1% and from the 9^th^ to 16^th^ day (*w*_*9-16*_) – 27.4 ± 27.0% (Figure 2). Moreover, in some rats of the group 1 the area of wounds increased – the *w* value was negative: minimum *w_*4-9*_* was −16.7%, maximum *w*_*4-9*_ was 40%. Minimum and maximum *w*_*9-16*_ were −33.3% and 60%, respectively.

**Fig. 2.**
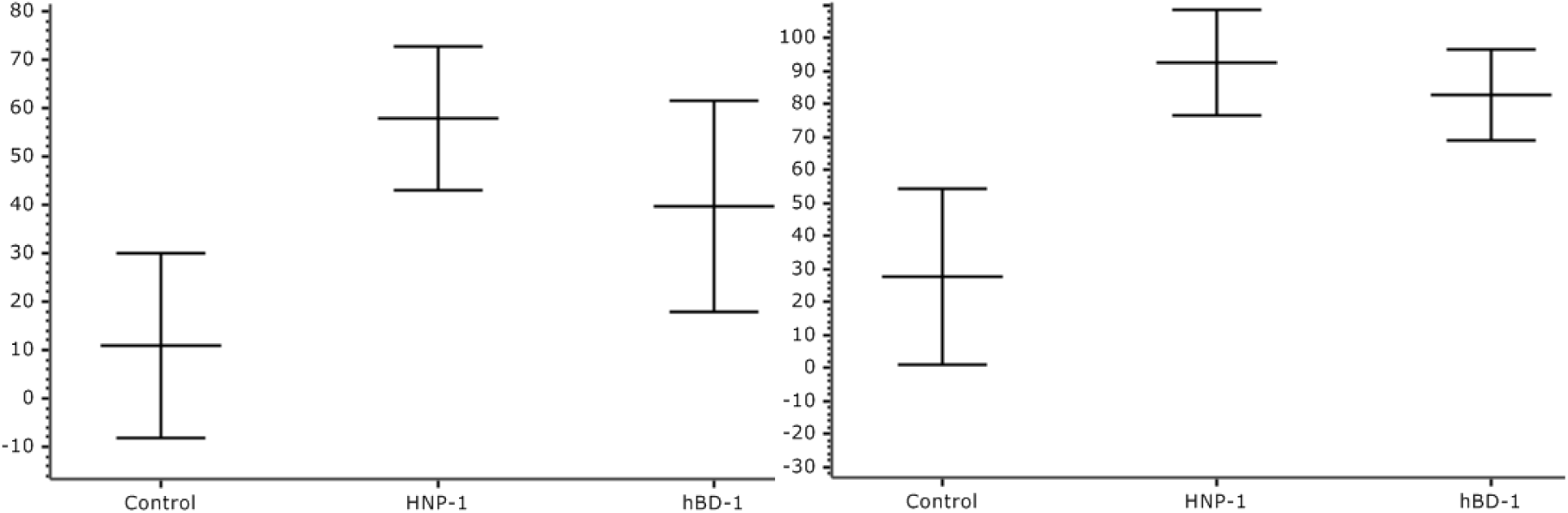
Wound healing rate (%, ±SD) from 4^th^ to 9^th^ day (left graph) and from 9^th^ to 16^th^ day (right graph) of the experiment.

Rats treated with niosomal HNP-1 gel (group 2) showed a high healing rate of infected wounds compared to control: *w*_*4-9*_ was 57.7 ± 14.7% (p<0.001) and *w*_*9-16*_ was 92.7 ± 15.8% (p<0.001).

Similar results were obtained in group 3 (niosomal gel with 1 mg/L of hBD-1): wound regeneration from the 4^th^ to 9^th^ day of the experiment was 39.7 ± 21.8% (statistically significant difference from the control (p<0.01), no difference from group 2) and from the 9^th^ to 16^th^ day – 82.8 ± 13.8% (statistically significant difference compared to control (p<0.001), no difference compared to group 2; Figure 2). Tables with wound areas and regeneration rates of each rat are available as Supplementary data.

## DISCUSSION

Our study showed that recombinant antimicrobial peptides, defensins HNP-1 and hBD-1, have a pronounced microbicidal activity against both MSSA and MRSA clinical strains. Moreover, these peptides were equally active against both strains – the MIC value for α-defensin-1 was 1 mg/L and that for β-defensin-1 was 0.5 mg/L. The lack of difference in the values of MIC for MSSA and MRSA can be explained by the fact that defensins do not affect penicillin-binding proteins. On the other hand, it would be interesting to conduct a study of protein-protein interactions in pairs of antimicrobial peptide-bacterial surface protein. This may help discover new mechanisms of AMP’s action. For example, it has been shown that α-defensin-1 can bind to cell wall precursor lipid II and inhibit cell wall synthesis [19].

The antimicrobial effect of the combination of cefotaxime with HNP-1/hBD-1 on MSSA is summed up. However, according to calculated fractional inhibitory concentration index, there was no synergy effect between HNP-1/hBD-1 + cefotaxime and the same was for the pair of HNP-1 + hBD-1. The latter can be explained by the same mechanism of antimicrobial action of α-defensin-1 and β-defensin-1. As for MRSA, neither synergy nor antagonism effects were observed since this strain has natural resistance to cefotaxime.

However, from our point of view, defensins cannot be considered as only antimicrobials. Defensins are modulators of inflammation and activators of the adaptive immune response. These antimicrobial peptides directly stimulate the migration of immune cells, promote the release of pro-inflammatory cytokines and activate antigen-presenting cells in the direction of the Th1-immune response thus, defensins are an effective link between innate and adaptive immunity [5].

As mentioned earlier, the practical use of antimicrobial peptides is complicated by their rapid degradation. To overcome this, we proposed to encapsulate AMPs in silica nanoparticles (niosomes). The results of *in vivo* experiments showed that topical gels with niosomal HNP-1/hBD-1 increase the healing rate of MRSA-infected wounds, compared to the control. In our previous study, we also demonstrated that niosomal hBD-1 increases the regeneration rate of MSSA-infected wounds in rats (a similar protocol was used, the duration of treatment was 13 days) [17]. Infected wound is a simple model of an infectious process, which helped us show that defensins work *in vivo* and have good prospects for clinical use. Therefore, it would be interesting to investigate parenteral or oral niosomal AMPs in models of generalized infection. It should be noted that defensin-mediated wound healing in this study may be associated not only with the antistaphylococcal activity of HNP-1/hBD-1. α- and β-defensins have been previously shown to have a chemotactic and wound healing effect [20–22]. β-defensin-1 is able to stimulate neutrophils to release so-called extracellular traps that “capture” *S. aureus*; moreover, this protective mechanism is triggered by stimuli from staphylococcal toxin [23]. Also, β-defensins induce pro-inflammatory cytokine production, cell migration and vasculogenesis in a wound [24].

## CONCLUSIONS

In the first part of the study, we showed for the first time the antimicrobial activity of recombinant defensins HNP-1 and hBD-1 against clinical strains of *S. aureus* (MSSA and MRSA) isolated from hospitalized patients with infected diabetic foot. In addition, we showed that these antimicrobial peptides do not have a synergistic effect with the β-lactam antibiotic cefotaxime against MSSA. β-defensin-1 had a higher anti-MSSA and anti-MRSA activity compared with α-defensin-1. Both AMPs had a lower MIC compared with cefotaxime (for MSSA). The results obtained *in vitro* using checkerboard method allowed us to choose the optimal concentrations of defensins for the preparation of topical gels with HNP-1/hBD-1 encapsulated in silica nanoparticles. The obtained antimicrobial gels were investigated in an experimental MRSA-infected wound model in rats. Gels containing HNP-1 and hBD-1 showed a significantly faster wound healing in comparison with the control. This study opens up new strategies for the development of new effective antibiotics based on antimicrobial peptides.

## ACKNOWLEDGEMENTS

This work was funded by Russian Foundation for Basic Research according to the research project No. 18-315-00081\18

## REFERENCES

[1] Iwasaki A., Medzhitov R. Control of adaptive immunity by the innate immune system. Nat. Immunol. (2015) 16 343–353.

[2] Martínez J.L., Baquero F. Emergence and spread of antibiotic resistance: Setting a parameter space. Ups. J. Med. Sci. (2014) 119 68–77.

[3] Martinez J.L., Fajardo A., Garmendia L. et al. A global view of antibiotic resistance. FEMS Microbiol. Rev. (2009) 33 44–65.

[4] Pachón-Ibáñez M.E., Smani Y., Pachón J. et al. Perspectives for clinical use of engineered human host defense antimicrobial peptides. FEMS Microbiol. Rev. (2017) 41 323–342.

[5] Suarez-Carmona M., Hubert P., Delvenne P. et al. Defensins: “Simple” antimicrobial peptides or broad-spectrum molecules? Cytokine Growth Factor Rev. (2015) 26 361–370.

[6] Steckbeck J.D., Deslouches B., Montelaro R. C. Antimicrobial peptides: new drugs for bad bugs? Expert. Opin. Biol. Ther. (2014) 14 11–14.

[7] Watermann A., Brieger J., Watermann A. et al. Mesoporous Silica Nanoparticles as Drug Delivery Vehicles in Cancer. Nanomaterials (2017) 7 pii=e189.

[8] Tong S.Y.C., Davis J.S., Eichenberger E. et al. Staphylococcus aureus infections: Epidemiology, pathophysiology, clinical manifestations, and management. Clin. Microbiol. Rev. (2015) 28 603–661.

[9] Matuschek E., Brown D.F.J., Kahlmeter G. Development of the EUCAST disk diffusion antimicrobial susceptibility testing method and its implementation in routine microbiology laboratories. Clin. Microbiol. Infect. (2014) 20 O255–266.

[10] White R.L., Burgess D.S., Manduru M. et al. Comparison of three different in vitro methods of detecting synergy: Time-kill, checkerboard, and E test. Antimicrob. Agents Chemother. (1996) 40 1914–1918.

[11] Orhan G., Bayram A., Zer Y. et al. Synergy tests by E test and checkerboard methods of antimicrobial combinations against Brucella melitensis. J. Clin. Microbiol. (2005) 43 140–143.

[12] Leverstein-van Hall M.A., Waar K., Muilwijk J. et al. Consequences of switching from a fixed 2:1 ratio of amoxicillin/clavulanate (CLSI) to a fixed concentration of clavulanate (EUCAST) for susceptibility testing of Escherichia coli. J. Antimicrob. Chemother. (2013) 68 2636–2640.

[13] Pfaller M.A., Espinel-Ingroff A., Boyken L. et al. Comparison of the broth microdilution (BMD) method of the European Committee on Antimicrobial Susceptibility Testing with the 24-hour CLSI BMD method for testing susceptibility of Candida species to fluconazole, posaconazole, and voriconazole by use of ep. J. Clin. Microbiol. (2011) 49 845–850.

[14] Milly P.J., Toledo R.T., Ramakrishnan S. Determination of minimum inhibitory concentration of liquid smoke fractions. J. Food. Sci. (2005) 70 5–16.

[15] Ruden S., Hilpert K., Berditsch M. et al. Synergistic interaction between silver nanoparticles and membrane-permeabilizing antimicrobial peptides. Antimicrob. Agents Chemother. (2009) 53 3538–3540.

[16] Odds F.C. Synergy, antagonism, and what the chequerboard puts between them. J. Antimicrob. Chemother. (2003) 52 1–1.

[17] Bolatchiev A., Baturin V., Bazikov I. et al. Effect of niosomal antimicrobial peptide hBD-1 on the healing rate of infected wounds in rats. Med. News North Caucasus (2018) 13 515–517.

[18] Zhao Y., Wang Z., Zhang Q. et al. Accelerated skin wound healing by soy protein isolate–modified hydroxypropyl chitosan composite films. Int. J. Biol. Macromol. (2018) 118 1293–1302.

[19] Leeuw E. de, Li C., Zeng P. et al. Functional interaction of human neutrophil peptide-1 with the cell wall precursor lipid II. FEBS Lett. (2010) 584 1543–1548.

[20] Grigat J., Soruri A., Forssmann U. et al. Chemoattraction of Macrophages, T Lymphocytes, and Mast Cells Is Evolutionarily Conserved within the Human α-Defensin Family. J. Immunol. (2007) 179 3958–3965.

[21] Funderburg N., Lederman M.M., Feng Z. et al. Human β-defensin-3 activates professional antigen-presenting cells via Toll-like receptors 1 and 2. Proc. Natl. Acad. Sci. (2007) 104 18631–18635.

[22] Boniotto M., Jordan W.J., Eskdale J. et al. Human β-defensin 2 induces a vigorous cytokine response in peripheral blood mononuclear cells. Antimicrob. Agents Chemother. (2006) 50 1433–1441.

[23] Kraemer B.F., Campbell R.A., Schwertz H. et al. Novel anti-bacterial activities of β-defensin 1 in human platelets: Suppression of pathogen growth and signaling of neutrophil extracellular trap formation. PLoS Pathog. (2011) 7 pii=e1002355.

[24] Roupé K.M., Nybo M., Sjöbring U. et al. Injury is a major inducer of epidermal innate immune responses during wound healing. J. Invest. Dermatol. (2010) 130 1167–1177.

