## Supplementary data for "Effect of Antimicrobial Peptides HNP-1 and hBD-1 on *Staphylococcus aureus* Clinical Strains *in vitro* and *in vivo*"

**Pages 1-5 – Synergy tests by chequerboard method results**
**Pages 6-7 – Wound healing rate results.**

### *Synergy tests by chequerboard method results.*

A – HNP-1 + cefotaxime against MSSA

B – hBD-1 + cefotaxime against MSSA

C – HNP-1 + hBD-1 against MSSA

D – HNP-1 + cefotaxime against MRSA

E – hBD-1 + cefotaxime against MRSA

F – HNP-1 + hBD-1 against MRSA

White cells – no growth.

Gray cells – visual presence of colony growth.

C-1 – control-1 – only 2.1% Mueller-Hinton broth (200 µl per well, without bacteria and without antimicrobial agents).

C-2 – control-2 – only bacterial inoculum (200 µl per well, without antimicrobial agents).

C-3 – control-3 – only antimicrobial agents without inoculum (100 µl of substance A + 100 µl of substance B) at maximal concentrations.

| | | | | | | | | | | | |
|---|---|---|---|---|---|---|---|---|---|---|---|
| **A** | **MSSA** | | | | | | | | | | |
| | **C-1** | **C-1** | **C-1** | **C-2** | **C-2** | **C-2** | **C-3** | **C-3** | **C-3** | | |
| **HNP-1 (mg/L)** | **5** | | | | | | | | | | |
| | **2.5** | | | | | | | | | | |
| | **1** | | | | | | | | | | |
| | **0.5** | | | | | | | | | | |
| | **0.25** | | | | | | | | | | |
| | **0.125** | | | | | | | | | | |
| | **0** | | | | | | | | | | |
| | | **0** | **0.125** | **0.25** | **0.5** | **1** | **2** | **4** | **8** | **16** | **32** |
| | **Cefotaxime (mg/L)** | | | | | | | | | | |

| | | | | | | | | | | | |
|---|---|---|---|---|---|---|---|---|---|---|---|
| **B** | **MSSA** | | | | | | | | | | |
| | **C-1** | **C-1** | **C-1** | **C-2** | **C-2** | **C-2** | **C-3** | **C-3** | **C-3** | | |
| **hBD-1 (mg/L)** | **5** | | | | | | | | | | |
| | **2.5** | | | | | | | | | | |
| | **1** | | | | | | | | | | |
| | **0.5** | | | | | | | | | | |
| | **0.25** | | | | | | | | | | |
| | **0.125** | | | | | | | | | | |
| | **0** | | | | | | | | | | |
| | | **0** | **0.125** | **0.25** | **0.5** | **1** | **2** | **4** | **8** | **16** | **32** |
| | **Cefotaxime (mg/L)** | | | | | | | | | | |

| C | MSSA | | | | | | | | |
|---|---|---|---|---|---|---|---|---|---|
| | C-1 | C-1 | C-1 | C-2 | C-2 | C-2 | C-3 | C-3 | C-3 |
| HNP-1 (mg/L) | 5 | | | | | | | | |
| | 2.5 | | | | | | | | |
| | 1 | | | | | | | | |
| | 0.5 | | | | | | | | |
| | 0.25 | | | | | | | | |
| | 0.125 | | | | | | | | |
| | 0 | | | | | | | | |
| | | 0 | 0.125 | 0.25 | 0.5 | 1 | 2.5 | 5 | |
| | hBD-1 (mg/L) | | | | | | | | |

| D | MRSA | | | | | | | | | | |
|---|---|---|---|---|---|---|---|---|---|---|---|
| | C-1 | C-1 | C-1 | C-2 | C-2 | C-2 | C-3 | C-3 | C-3 | | |
| HNP-1 (mg/L) | 5 | | | | | | | | | | |
| | 2.5 | | | | | | | | | | |
| | 1 | | | | | | | | | | |
| | 0.5 | | | | | | | | | | |
| | 0.25 | | | | | | | | | | |
| | 0.125 | | | | | | | | | | |
| | 0 | | | | | | | | | | |
| | | 0 | 0.125 | 0.25 | 0.5 | 1 | 2 | 4 | 8 | 16 | 32 |
| | Cefotaxime (mg/L) | | | | | | | | | | |

| E | MRSA | | | | | | | | | |
|---|---|---|---|---|---|---|---|---|---|---|
| | C-1 | C-1 | C-1 | C-2 | C-2 | C-2 | C-3 | C-3 | C-3 | |
| hBD-1 (mg/L) | 5 | | | | | | | | | | |
| | 2.5 | | | | | | | | | | |
| | 1 | | | | | | | | | | |
| | 0.5 | | | | | | | | | | |
| | 0.25 | | | | | | | | | | |
| | 0.125 | | | | | | | | | | |
| | 0 | | | | | | | | | | |
| | | 0 | 0.125 | 0.25 | 0.5 | 1 | 2 | 4 | 8 | 16 | 32 |
| | Cefotaxime (mg/L) | | | | | | | | | | |

| F | MRSA | | | | | | | | |
|---|---|---|---|---|---|---|---|---|---|
| | C-1 | C-1 | C-1 | C-2 | C-2 | C-2 | C-3 | C-3 | C-3 |
| HNP-1 (mg/L) | 5 | | | | | | | | |
| | 2.5 | | | | | | | | |
| | 1 | | | | | | | | |
| | 0.5 | | | | | | | | |
| | 0.25 | | | | | | | | |
| | 0.125 | | | | | | | | |
| | 0 | | | | | | | | |
| | | 0 | 0.125 | 0.25 | 0.5 | 1 | 2.5 | 5 | |
| | hBD-1 (mg/L) | | | | | | | | |

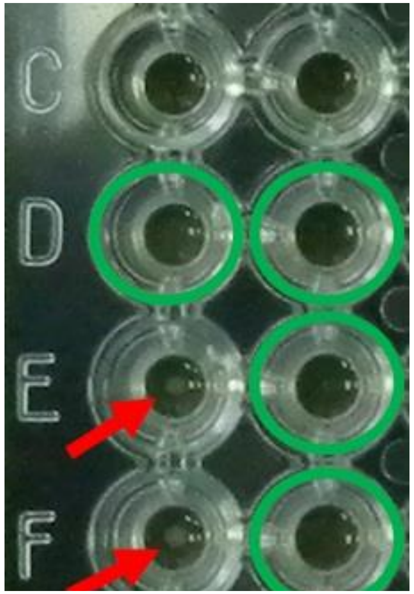


Example of visual assessment of bacterial growth after incubation with antimicrobial agents using the chequerboard method: the presence of the growth of *S. aureus* colonies is indicated by red arrows, the absence of growth is indicated by green circles.

***Effect of HNP-1/hBD-1 encapsulated in silica nanoparticles on MRSA-infected wound healing in rats.***

Wound area (S) and wound perimeter (P) for each animal in all groups.

| | **Group 1 (control)** | | | | | | **HNP-1, 2 mg/L** | | | | | | **hBD-1, 1 mg/L** | | | | | |
|---|---|---|---|---|---|---|---|---|---|---|---|---|---|---|---|---|---|---|
| Rat | **Day 4** | | **Day 9** | | **Day 16** | | **Day 4** | | **Day 9** | | **Day 16** | | **Day 4** | | **Day 9** | | **Day 16** | |
| | S, $mm^2$ | P, mm | S, $mm^2$ | P, mm | S, $mm^2$ | P, mm | S, $mm^2$ | P, mm | S, $mm^2$ | P, mm | S, $mm^2$ | P, mm | S, $mm^2$ | P, mm | S, $mm^2$ | P, mm | S, $mm^2$ | P, mm |
| 1 | 5 | 22 | 5 | 25 | 3 | 16 | 4 | 25 | 1 | 13 | 0 | 0 | 5 | 27 | 4 | 25 | 1 | 15 |
| 2 | 6 | 30 | 4 | 19 | 3 | 19 | 9 | 40 | 5 | 29 | 4 | 26 | 6 | 31 | 4 | 25 | 0 | 0 |
| 3 | 5 | 27 | 3 | 18 | 3 | 19 | 5 | 29 | 2 | 15 | 0 | 0 | 6 | 27 | 5 | 26 | 0 | 0 |
| 4 | 6 | 26 | 6 | 29 | 5 | 25 | 7 | 38 | 4 | 24 | 2 | 17 | 8 | 33 | 5 | 26 | 3 | 25 |
| 5 | 5 | 28 | 4 | 25 | 2 | 18 | 5 | 27 | 2 | 18 | 0 | 0 | 7 | 32 | 6 | 28 | 2 | 18 |
| 6 | 6 | 25 | 6 | 30 | 5 | 28 | 8 | 36 | 2 | 16 | 0 | 0 | 7 | 30 | 3 | 20 | 2 | 17 |
| 7 | 5 | 25 | 5 | 26 | 4 | 20 | 5 | 27 | 1 | 14 | 0 | 0 | 7 | 31 | 4 | 24 | 1 | 14 |
| 8 | 6 | 28 | 4 | 19 | 3 | 20 | 6 | 31 | 3 | 19 | 0 | 0 | 4 | 24 | 1 | 13 | 0 | 0 |
| 9 | 7 | 35 | 7 | 32 | 6 | 32 | 4 | 24 | 2 | 18 | 0 | 0 | 7 | 32 | 2 | 20 | 2 | 19 |
| 10 | 6 | 27 | 7 | 31 | 8 | 34 | 5 | 29 | 3 | 21 | 0 | 0 | 7 | 38 | 5 | 29 | 1 | 10 |

Wound healing rate (%) for each animal, calculated from 4th to 9th day of the experiment ($w_{4\text{-}9}$) and from 9th to 16th day of the experiment ($w_{9\text{-}16}$).

| Rat | Group 1 (control) | | HNP-1, 2 mg/L | | hBD-1, 1 mg/L | |
|---|---|---|---|---|---|---|
| | $w_{4\text{-}9}$ | $w_{9\text{-}16}$ | $w_{4\text{-}9}$ | $w_{9\text{-}16}$ | $w_{4\text{-}9}$ | $w_{9\text{-}16}$ |
| **1** | 0.0 | 40.0 | 75.0 | 100.0 | 20.0 | 80.0 |
| **2** | 33.3 | 50.0 | 44.4 | 55.6 | 33.3 | 100.0 |
| **3** | 40.0 | 40.0 | 60.0 | 100.0 | 16.7 | 100.0 |
| **4** | 0.0 | 16.7 | 42.9 | 71.4 | 37.5 | 62.5 |
| **5** | 20.0 | 60.0 | 60.0 | 100.0 | 14.3 | 71.4 |
| **6** | 0.0 | 16.7 | 75.0 | 100.0 | 57.1 | 71.4 |
| **7** | 0.0 | 20.0 | 80.0 | 100.0 | 42.9 | 85.7 |
| **8** | 33.3 | 50.0 | 50.0 | 100.0 | 75.0 | 100.0 |
| **9** | 0.0 | 14.3 | 50.0 | 100.0 | 71.4 | 71.4 |
| **10** | -16.7 | -33.3 | 40.0 | 100.0 | 28.6 | 85.7 |
